# *In vivo* single-cell analysis using calcofluor - white staining detects high expression phenotype in *L. lactis* cultures engineered for hyaluronic acid production

**DOI:** 10.1101/2020.10.21.348672

**Authors:** Anantha-Barathi Muthukrishnan, Antti Häkkinen, Velvizhi Devi Rajendran, Anupama Kozhiyalam, Guhan Jayaraman

## Abstract

Hyaluronic acid (HA) is a biopolymer with wide applications in the field of medicine and cosmetics. Bacterial production of HA has a huge market globally. Certain species of *Streptococcus* are native producers of HA but they are pathogenic. Therefore, safer organisms such as *L. lactis* are engineered for HA production. However, there are challenges such as low yield, low molecular weight and polydispersity of HA obtained from these cultures. Optimisation of bioprocess parameters and downstream purification parameters are being addressed to overcome these challenges. We explore these problems from the perspective of microbial heterogeneity, since variations in phenotype affect the yield and properties of the product in a bioreactor. For this perspective, a method to quantitatively assess the occurrence of heterogenous phenotypes depending on the amount of HA produced at the single-cell level is required. Here, we evaluated for the first time the use of calcofluor white staining method combined with *in vivo* fluorescence confocal microscopy to quantify the heterogeneity in phenotypes of *L. lactis* cells engineered for HA production.

From the microscopy image analysis, we found that the population harbours significant heterogeneity with respect to HA production and our novel approach successfully differentiates these phenotypes. Using the fluorescence intensity levels, first we were able to confidently differentiate cells not expressing HA (Host cells without HA genes for expression) from cells with genes for HA production (GJP2) and induced for expression, as there is a consistently two-fold higher level of expression in the GJP2 cells independently of the cell size. Further, this method revealed the occurrence of two different phenotypes in GJP2 cultures, one of a high-expression phenotype (40% of the population) and the other one of a low-expression (remaining 60% of the population), and it is the high expression phenotype that contributes to the increase in the HA expression of the GJP2 population compared with the host cells. Thus, it is essential to identify the extrinsic and intrinsic factors that can favour most of the cells in the population to switch and stabilise into the high-expression phenotype state in a bioreactor, for higher yield and possibly reduced heterogeneity of the product, such as polydispersity in chain lengths. For such optimisation studies, this *in vivo* method serves as a promising tool for rapid detection of phenotypes in the bioreactor samples under varying conditions, allowing fine tuning of the factors to stabilise high-expression phenotypes thereby maximizing the yield.

**Graphical Abstract:** done.

**Key Points:**

- Calcofluor staining successfully differentiated the phenotypes based on HA levels.
- This study revealed the occurrence of significant heterogeneity in HA expression.
- This method will aid for rapid optimization of factors for improved HA production.

## INTRODUCTION

Hyaluronic acid (HA) is a natural biopolymer with wide applications for cosmetics and medicine industry. High molecular weight HA formulations applied topically over skin promote wound healing (Kogan *et al.*, 2006). The antioxidant properties of HA can be used as anti-inflammatory component in wound dressing material (Moseley *et al.*, 2003). HA is a linear mucopolysaccharide consisting repetitive units of glucuronic acid and N-acetyl glucosamine linked by β 1-3 and β 1-4 glycosidic bonds (**Figure 1A).** HA is initially obtained from animal sources and later from microbes, due to various reasons including the proteoglycans associated with rooster combs, which makes the isolation process tedious and costly (Chien *et al.*, 2007).

**Figure 1:**
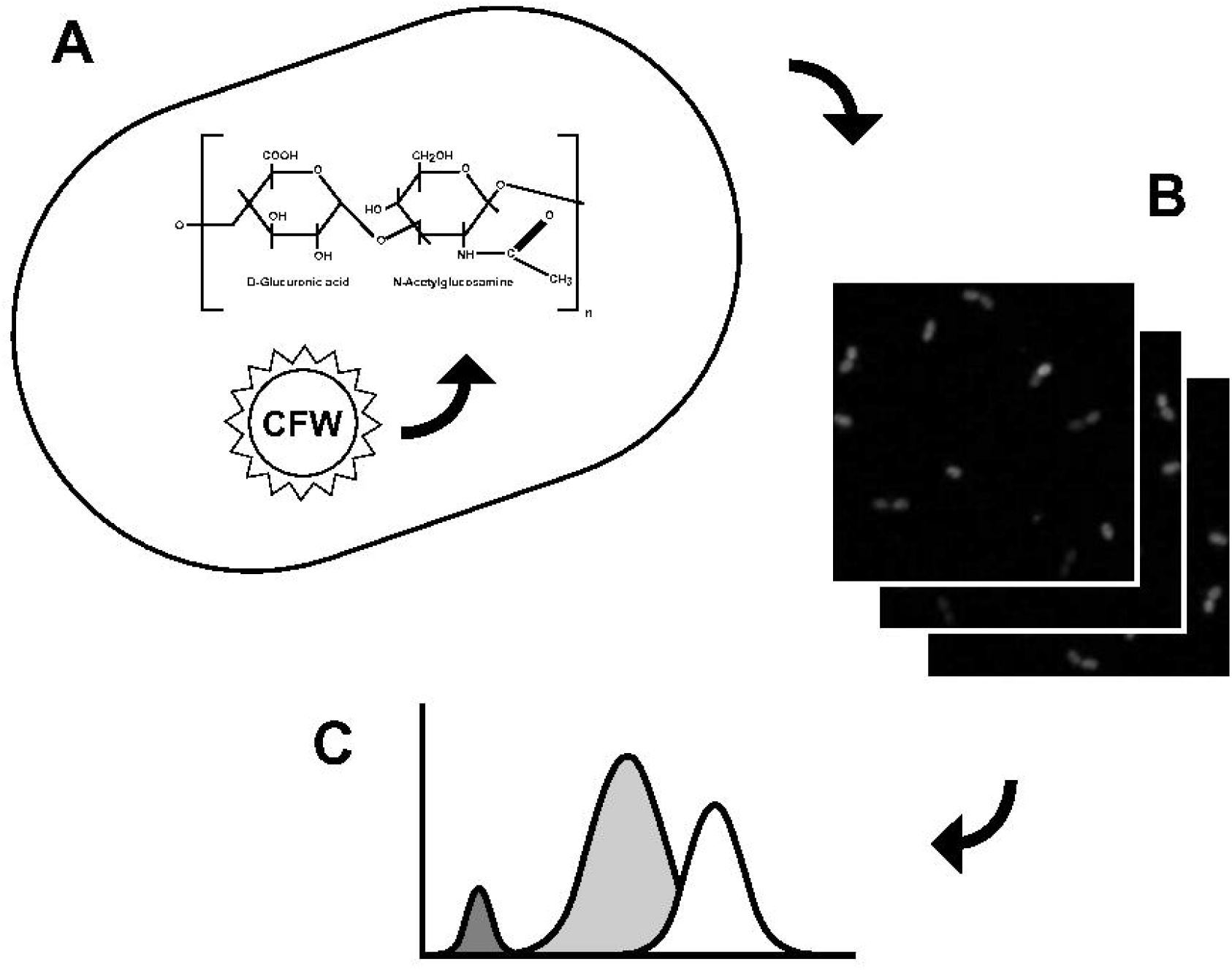
Schematic illustration of the proposed in vivo monitoring strategy of the HA producing *L. lactis* cells. (A) The strains were engineered produces hyaluronic acid (HA), which is bound by the calcofluor stain (CFW). This process can be monitored in single live cells under a fluorescence microscope. (B) Microscope images of a normal *L. lactis* and the HA producing strain were acquired and subjected to image analysis. (C) A statistical analysis of these data suggests three phenotypes: dark and low expressing cells, which are present in the normal phenotype, and a high expression phenotype, which coexists with the low expression phenotype in the engineered strain only, and is responsible for a significant HA production.

*Streptococcus* is the widely used bacterial source of HA today but it poses serious threats such as bacterial endotoxins, which makes the downstream purification of *streptococcal* fermented HA cost ineffective (Liu *et al.*, 2011). For this reason, several attempts have been made to produce HA from organisms generally regarded as safe. We engineered a heterologous HA synthesis pathway in *Lactococcus lactis,* a GRAS organism, which is employed widely in the food industry for the production and preservation of fermented products (Chien *et al.*, 2007). The recombinant bacterial hosts for HA are successfully producing HA. However when compared with the yield of the native producers such as the *S. zooepidemicus* produces 6-7 g/L, the yield is much lower (**Table 2**).

Further, another major challenge in HA production is polydispersity, the occurrence of HA polymer as a mixture composed of non-uniform chain lengths. This is highly disadvantageous, as the HA chain length or molecular weight determines the specific properties of HA necessary for product formulation. For example, shorter chain lengths are ideal for cosmetics, medium chain lengths are capable to address the pro angiogenic and anti-apoptotic properties that stimulates the synthesis of Heat Shock Proteins (HSP) and are potent immunostimulants. The very long chain lengths are suitable for medical applications such as immunosuppression and anti-angiogenic functions. (Natalia *et al.*, 2016) Thus, highly polydispersed HA needs sophisticated downstream processes to achieve uniform chain lengths, making HA a costly raw material. Thus, it is essential to address these two challenges, namely, yield and polydispersity, to make recombinant HA production a safer choice of source and to become an economically viable alternate for the current sources with potential risks (Cansu *et al.*, 2018).

Several factors have been addressed to improve the yield of HA produced by *L. lactis* cells such as increasing concentrations of HA precursors (Jeeva *et al.*, 2019), process optimization strategies (Puvendran *et al.*, 2019) and integration of HA pathway genes in the genome (Hmar *et al.*, 2014). Also, there are studies on polydispersity issues regarding the HA M_w_, which is high in the initial glucose concentrations and experiences a drop in a specific growth rate during glucose uptake (David *et al.*,1997), suggesting that polydispersity can be due to one of the major factors such as the precursor (UDP-GlcUA and UDP-GlcNAc) concentrations in the appropriate ratio. This is tunable by over expressing the HA pathway genes coding for the enzymes that catalyse the HA synthesis (Jeeva *et al.*, 2019).

In addition to these, microbial heterogeneity has been identified as one of the major factor affecting recombinant phenotypes and thereby product yield and properties. In a colony of cells, the single or an individual cell, getting sorted as a producer or non-producer is observed during growth, survival and their productivity in order to gain insights of the weaknesses of the population as a whole (Koutsoumanis *et al.*, 2017). The ineffective clonal population of cells can create a remarkable impact on the activity of industrial cultures (Muller *et al.*, 2010). Microbial phenotypic heterogeneity is a vital component of biological diversity especially under highly fluctuating stressful environments. Such disturbances caused due to metabolic heterogeneity in the microenvironment of pilot scale bioreactors are less well understood, as the lab scale cultivation will rarely face such issues. Hence it is required to design analytical tools and metabolic engineering strategies to account for metabolic variability in bioprocess conditions (Delvigne *et al.*, 2014).

Heterogeneity is referred as a noise or outcome of a noisy environment in the cell population deviating from a typical, average cell. Heterogeneity may be genetic or non-genetic in nature, the latter including both intrinsic or extrinsic factors, which correspond to variability of the internal cell state and its environment, respectively. Quantifying the heterogeneity in clonal populations and building analytical tools for the understanding of sources of heterogeneity in biological systems can improve our insights on optimisation of both extrinsic and intrinsic factors that could overall improve the homogeneity of the population. Hence it is required to design analytical tools and metabolic engineering strategies that manipulate the metabolic variability in bioprocess conditions (Delvigne *et al.*, 2014).

Following this, we focused in understanding the impact of microbial heterogeneity on HA yield and polydispersity of metabolically engineered *L. lactis* strain, a less explored area in this context. This study visualizes the *L. lactis* population at the single-cell level to verify our hypothesis on occurrence of population heterogeneity and its impact with respect to HA production in the engineered *L. lactis* cells.

To our knowledge such studies have not been carried out so far. Therefore, first we developed a visualization method for real-time live-cell imaging of *L. lactis* population at the single-cell level using fluorescence confocal microscopy. For this, we employed a stain, namely, calcofluor white, which is an optical brightener with a high affinity towards β1-4 glucans such as cellulose, chitin and other β-linked polysaccharides (Huges *et al.*, 1981). This stain is also used in identifying fungal structures, spores and other fruiting body (Jensen *et al.*, 1998). Until now, calcofluor selection was not used for visualizing HA cells to the best of our knowledge, and there are many studies where the dye is efficiently used in staining the cellulose and visualization of banana and mango cells (Rongkaumpan *et al.*, 2019).

Specifically, in this study we first created a novel method for *in vivo* monitoring of HA producing *L. lactis* cells. Next, we verified the ability of the stain to differentiate the various HA producing phenotypes using image analysis methodology. Finally, we discovered that the phenotypes in the population of engineered *L. lactis* cells can be partitioned into a low- and a high expression phenotype with respect to HA production. Furthermore, we characterized the degree of heterogeneity that manifests at the population level.

## MATERIALS and METHODS

### Chemicals

The modified GM17 (glucose-M17) medium components (yeast extract 5 g/L, brain heart infusion 5 g/L, MgSO_4_·7H_2_O 0.5 g/L, ascorbic acid 0.5 g/L, KH_2_PO_4_ 0.5 g/L, and K_2_HPO_4_ 1.5 g/L), glucose, antibiotics (Chloramphenicol) CTAB (Cetyl trimethyl ammonium bromide), and agarose for microscopy sample preparation were purchased from Himedia Laboratories (India). Chemicals such as sulphuric acid, isopropyl alcohol, sodium nitrate and potassium hydroxide were purchased from Merck (USA). Calcofluor white for staining the HA capsule and nisin for induction of the cells were purchased from Sigma-Aldrich, India.

### Bacterial strain, plasmid and growth conditions

The *L. lactis* strain NZ9000 (NIZO, Netherlands), a widely used GRAS organism, was used as the heterologous host for HA production in this study. This host with the pNZ8148 vector (NIZO, Netherlands) carrying the *hasA* and *hasB* genes of the operon for HA synthesis under the *Pnis* promoter is a previously engineered *L. lactis* system from our lab, referred as GJP2 strain (Jeeva *et al.*, 2019). The genes *hasA* and *hasB* are from *Streptococcus zooepidemicus*, a lancefield group C opportunistic pathogen and code for key HA pathway enzymes, namely, HA synthase and UDP-glucose 6-dehygrogenase, respectively. For routine cultures, the host cells and GJP2 cells were grown in GM17 medium with glucose (0.5%). The antibiotic chloramphenicol (10 μg/mL) is additionally supplemented to the GJP2 cells for plasmid propagation. The cultures were incubated at 30°C, under static conditions.

### Induction of genes for HA production by engineered *L. lactis* cells

The cells were incubated overnight (~16 hours), and from the overnight culture, cells were inoculated into fresh medium with appropriate antibiotic at 1:1000 dilution. Cultures were incubated until they reached the exponential phase (~0.6 OD_600nm_). The *Pnis* promoter, a native promoter for *L. lactis,* is inducible externally with nisin, a polycyclic antibacterial peptide (Kuipers *et al.*, 1998). To find the optimal nisin concentration for maximum induction, we induced the same host strain with the same plasmid but with a mCherry reporter protein instead of the *hasA* and *hasB* genes under the *Pnis* promoter with varying concentration (0.5 to 3 ng/mL) of nisin. These cultures are then incubated for three hours and from that 1 OD of cells were taken and centrifuged. The cell pellets were washed twice in PBS and then finally resuspended in 500 μL of PBS. From these 150 μL were added in three wells of the microplate. The mean fluorescence of the mCherry reporter proteins and also cells without any induction (control) were measured using a Tecan microplate fluorometer with appropriate excitation (587 nm) and emission (620 nm) wavelengths. All the conditions were measured in an analogous fashion and repeated with three biological triplicates.

Based on the results, as shown in the Results section, 2 ng/mL of nisin was selected for further studies to induce the target genes *hasA* and *hasB* under the Pnis promoter.

### HA production in bioreactors

Cells were grown in an unaerated bioreactor (Bioengineering, Switzerland) with a maximum capacity of 2.4 L. For this, the M17 medium components (for 1.2 L) were dissolved and sterilized *in situ.* The optimum growth conditions maintained were 30°C, 200 rpm and pH 7. Sterile glucose (0.5%) along with the chloramphenicol (10μg/mL) was injected aseptically into the medium. The inoculum culture for the fermentation was prepared as follows: First, from the glycerol stocks stored at −80 °C, a loop full of cells were inoculated to 5 mL of medium with antibiotic in a test tube and grown overnight at optimum growth conditions. Next, the overnight seed culture was used to inoculate 100 mL of medium with antibiotic and incubated until it reached 0.6OD. This culture was then used as inoculum for the bioreactor fermentation at a final concentration of 5%. Immediately after inoculation, the 0^th^ hour samples were collected followed with the sample collection in regular intervals until the glucose was exhausted. All the samples were stored in −20 °C for further quantitative analysis for biomass detection and quantification of HA concentration and HA molecular weight. All the bioreactor runs were performed twice, and further studies were carried out from these duplicates.

### Estimation and detection of HA

To verify HA production by the induced GJP2 cells, the CTAB method was employed. HA is a polyanionic polymer which gets precipitated by the addition of CTAB, which is an organic ammonium cation. This precipitation results in turbidity which is proportionate to the concentration of HA. This turbidity is measured using a spectrophotometer. For this analysis, 1 mL of the culture was treated with equal volume of 0.1% Sodium dodecyl sulphate (SDS) and mixed by vortexing and incubated at room temperature for 10 minutes. Then the samples were centrifuged and the supernatant was collected for CTAB analysis as described previously (Oueslati *et al.*, 2014). To the micro-well plate the supernatant is added, followed by 0.1M phosphate buffer. The plate is kept on a gel rocker for 15 - 20 seconds and then incubated at 37 °C for 10 - 15 minutes, along with the CTAB reagent (2.5 g CTAB in 100mL of 2% NaOH). Then the plate was removed and the pre-incubated CTAB was added to all the samples. These samples were again kept on a gel rocker for a few seconds for thorough mixing and further incubated at 37°C for 10 - 15 minutes. The OD of these samples and also the host cells (control) were measured at 600nm.

To further confirm the presence of HA and to detect the molecular weight of HA, Size Exclusion Chromatography (SEC) was performed as described earlier (Sreeja *et al.*, 2017). The GJP2 samples identified as positive for HA production using CTAB and SEC were used as samples for further *in vivo* imaging studies.

### Calcofluor staining and *in vivo* imaging

Cells were treated with equal volume of calcofluor stain and incubated for 2 minutes at RT. Then from these samples 0.7-1 μL of the sample was added to the slide contacting 1% agarose gel and covered with a coverslip. Cells were imaged using a Zeiss LSM 700 (Germany) inverted confocal fluorescence laser scanning microscope using a 100X objective. Zen software was used for image acquisition and single time point images were acquired using a 405 nm ion laser and PMT for detection of fluorescence from the calcofluor dye. Z-stack images were also taken and a range indicator is used to avoid saturation while imaging. Image and data analysis of these experiments were done using separate signal processing steps.

### Image and data analysis

#### (i) Segmentation

The cell segmentation was performed as follows. First, the difference between adjacent Z-planes were computed, in order to discard in homogeneities and artifacts, such as dust particle responses and distortions, which generate a constant response in the bright field image. Second, for each differential plane, the cell edge responses were reconstructed using Wiener deconvolution with a Laplacian-of-Gaussian filter with a standard deviation of 3 pixels (the value was determined experimentally, and corresponds to the apparent cell minor radius, see **Figure 5**) and signal-to-noise ratio of 100%. Afterwards, the reconstructed planes were averaged in order to capture cells lying in different Z-planes, and the field was divided into Watershed segments. The segments, whose intensity signal was more likely to be generated by a constant signal as opposed to a ball-tapered signal with a standard deviation of 3 pixels (matching the above), in least-squares sense, were discarded as background. Finally, the segments which were adjacent to the image border were discarded, in order to remove partial cells which could bias the size (area) estimates. The segmentation was manually checked and ambiguous segments, such as cells rotating about the different Z-planes were discarded.

#### (ii) Expression quantification

The expression was quantified from the fluorescence channel of the microscope images as follows. First, the nine (9) adjacent Z-planes were merged by taking the pixelwise maximum across the Z-stack, resulting in a univariate intensity field across the imaged plane, and allows capturing the data from the optimally focused plane regardless of in which of them the cell lies on. Second, the resulting intensity field was summed up (integrated) inside each cell, as determined by the segmentation, to estimate the abundance of fluorescence molecules inside each cell. The resulting numbers are proportional to both the (average) cell brightness and the cellular footprint (area). As the fluorescence is expected to be abundant and uniformly distributed inside the cells (see **Figure 5**, the numbers were normalized using the corresponding cell area in order to derive an estimate of the expression level. Expression value of 0 corresponds to no response on the sensor in any Z-plane, while a value of 1 corresponds to fully saturated sensor values at the optimally focused Z-plane.

#### (iii) Expression clustering

The cells were classified into “dark”, “low” and “high” expression phenotypes as follows. First, by observing the intensity distributions, each of the Host phenotype images were assumed to contain the dark and low expression phenotypes, while the GJP2 phenotype images were assumed to contain the low and high expression phenotypes. The expression levels of the dark, low and high phenotypes are assumed to be ordered increasingly. By assuming each distribution can be approximated with a normal distribution with an unknown mean and unknown shared variance, as estimate minimizing the likelihood of generating the samples can formed (as in Otsu’s method (Otsu. 1979)). Specifically:

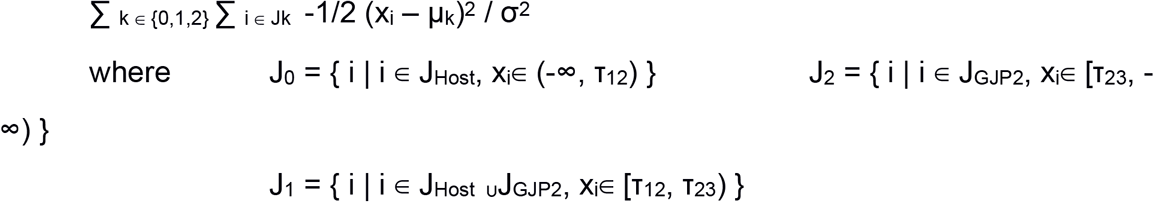

is maximized, where x_i_ is the expression level of the i:th sample; J_Host_ and J_GJP2_ the indices of the samples corresponding to the Host and GJP2 phenotypes, respectively; J_0_, J_1_, and J_2_ the unknown indices of the dark, low, and high expression samples, respectively; τ_12_ and τ_23_ the unknown thresholds between dark-low and low-high, respectively; μ_0_, μ_1_, and μ_2_ are unknown means of the dark, low, and high population, respectively, and σ^2^ is the unknown common intracluster variance. The global optimum can be found in O(n^2^) time using Otsu’s tabulation on (τ_12_, τ_23_) where n is the number of samples (Otsu, 1979).

## RESULTS

### Determination of HA production by engineered *L. lactis* cells

Our aim is to develop a visualisation method to observe HA production in engineered *L. lactis* cells *in vivo*. An overview of the procedure is shown in **Figure 1**. For this, we first identified the optimal concentration of the inducer (nisin) for HA production under the *Pnis* promoter by these cells using mCherry fluorescent reporter proteins. The cells were subjected to 0.5 to 3 ng/mL of nisin and were measured using a fluorometer. The host cells and the GJP2 cells without induction and with and without nisin were used as controls. Results from the measurements suggested that 2 ng/mL is the optimal concentration for higher expression (**Figure 2**, **Table 1**). Based on these results, this concentration is used for induction of the HA pathway genes *hasA* and *hasB*, which are kept under the same *Pnis* promoter of the same plasmid. Cell samples from bioreactor studies were analyzed for HA production as described in the methods section. From the CTAB analysis, the yield of HA was measured as 0.8g/L in the GJP2 cells and no production in the host cells, as expected (**Figure 3**, **Table 2**). Further, from SEC the HA peak was detected for the GJP2 cells and the molecular weight was found to be in the range of 0.2-1.2 MDa (**Figure 4**, **Table 3**). Thus, the production of HA by the GJP2 cells was confirmed and these samples were used for imaging using calcofluor staining.

**Figure 2:**
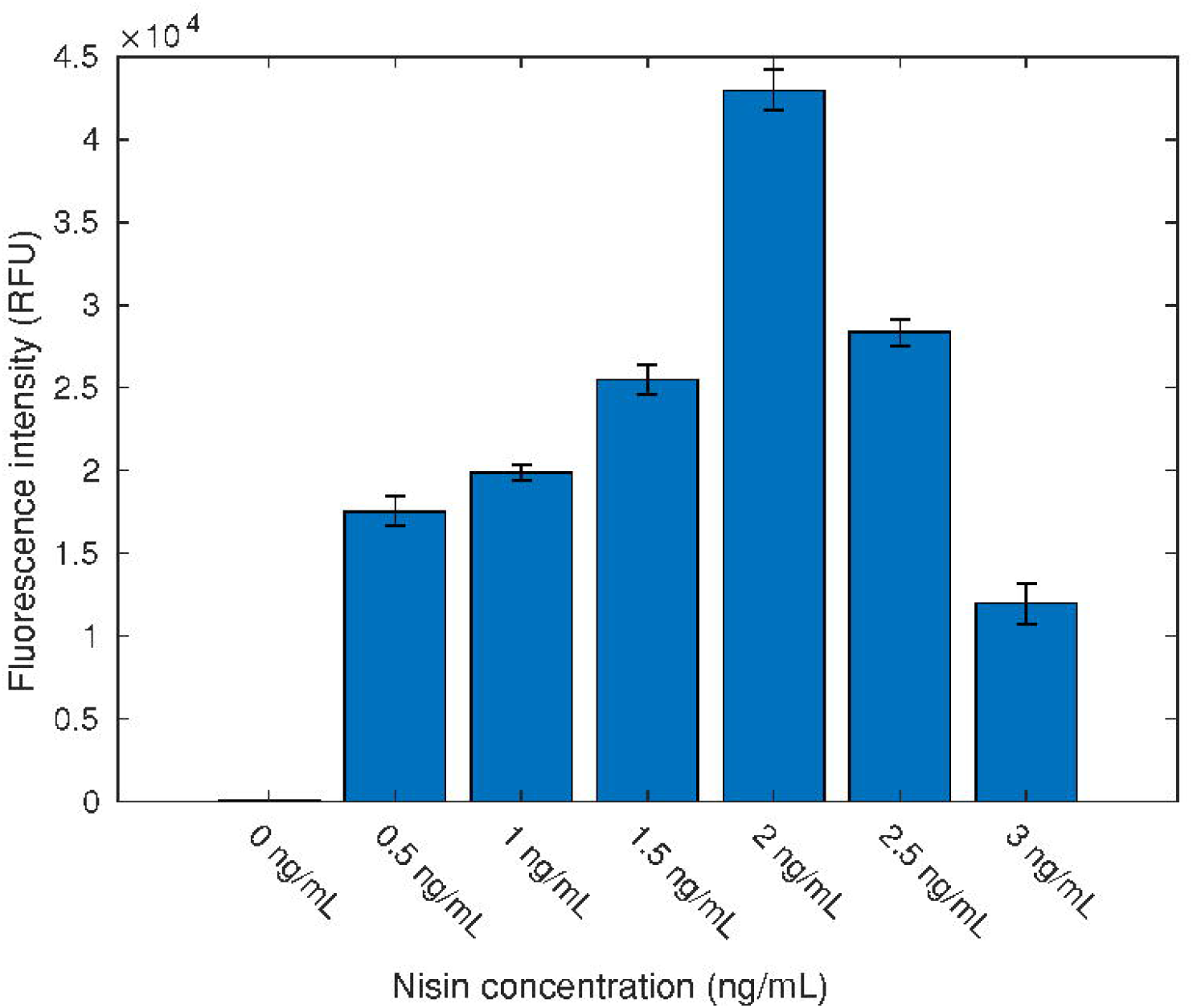
mCherry reporter production as quantified by the fluorometer as a function of the inducer (nisin) concentration. The bars denote averages from three replicates, and the whiskers one standard error of the mean. The data are tabulated in **Table 1**.

**Figure 3:**
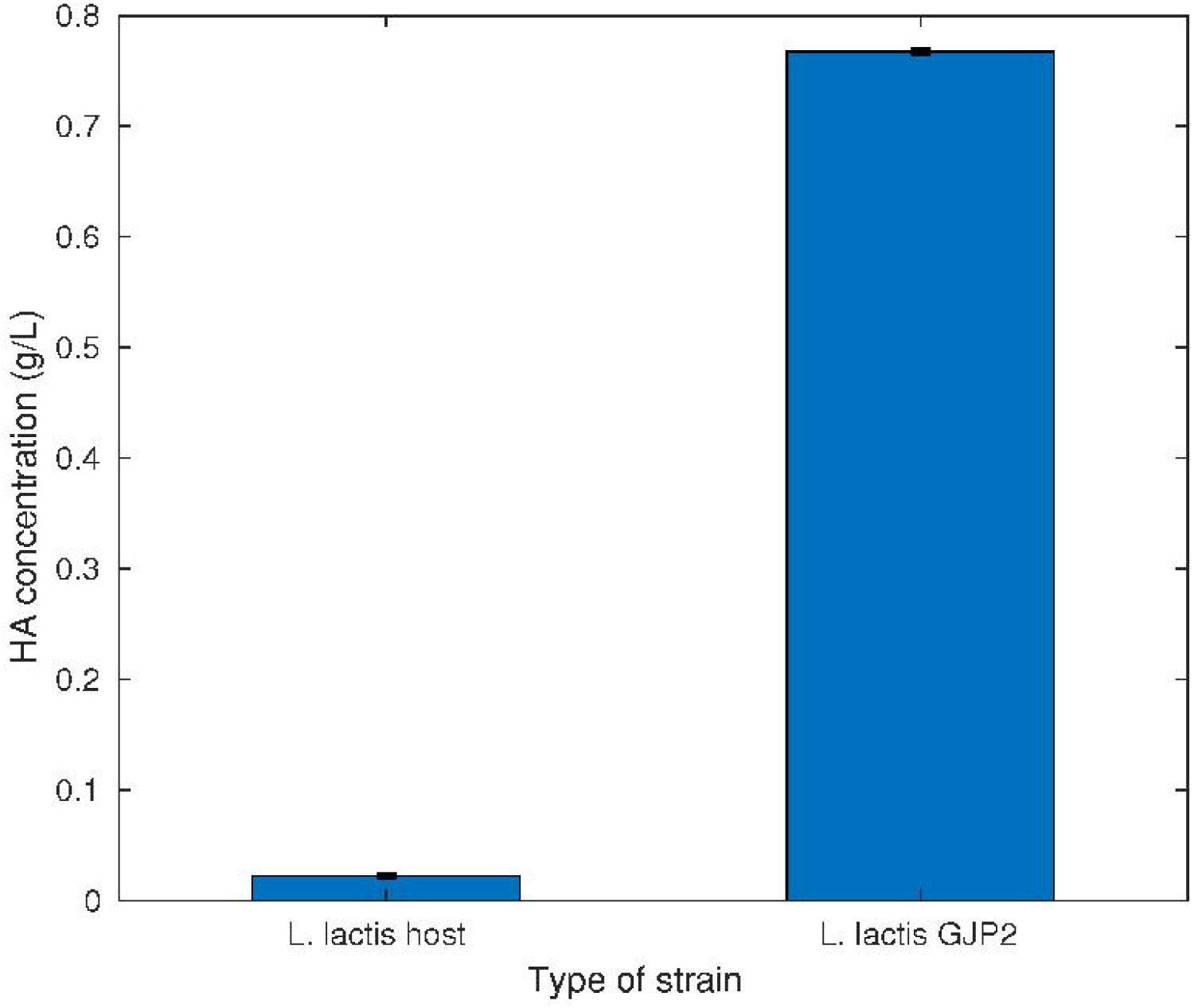
HA yield in bioreactor studies using the CTAB analysis. The bars denote averages from two replicates, and the whiskers one standard error of the mean. The data are tabulated in **Table 2**.

**Figure 4:**
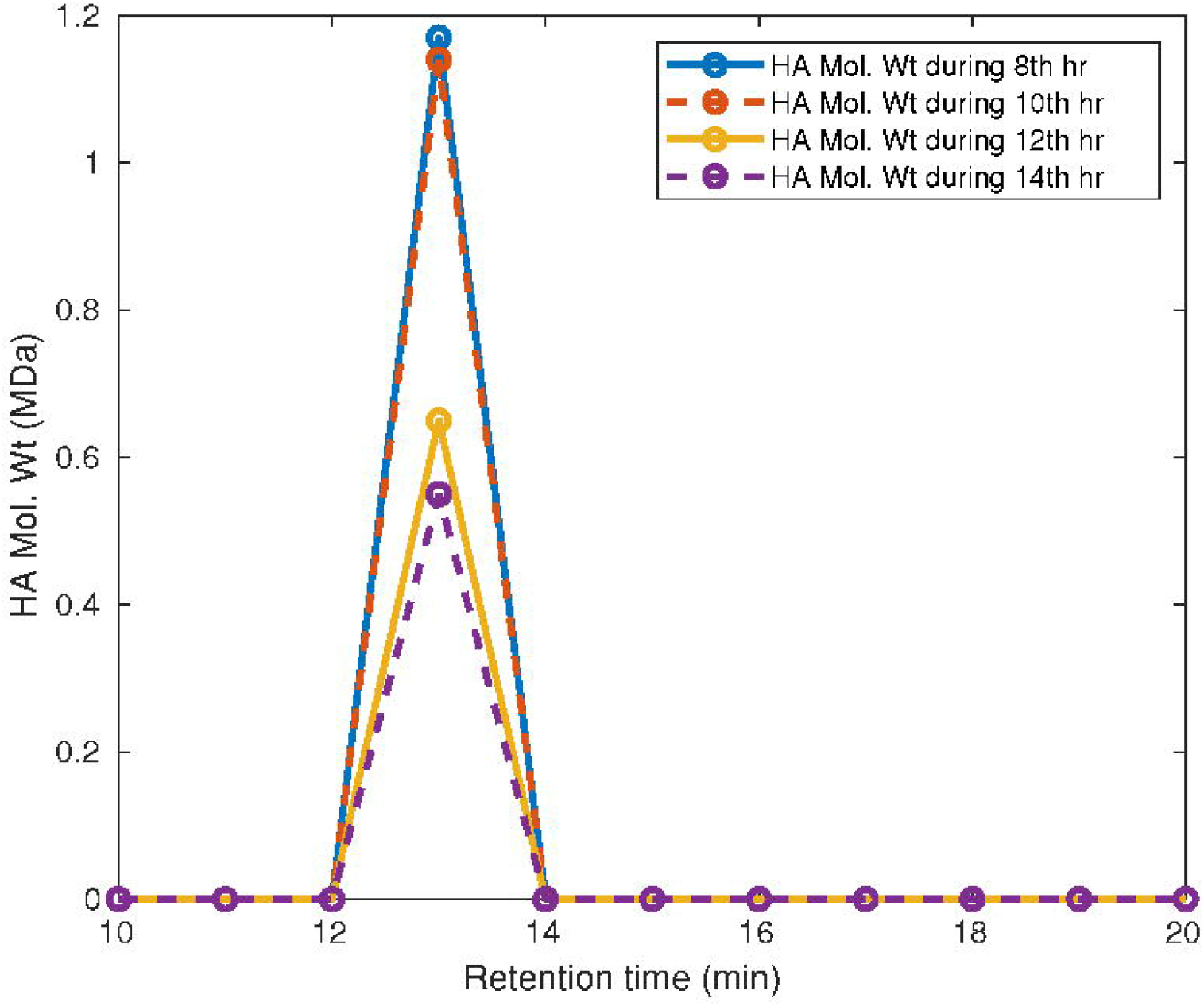
Molecular weight of the HA from size exclusion chromatography measurements. The circles represent measurements. The data are tabulated in **Table 3**.

### Calcofluor staining

The calcofluor stained cells were then visualized using a fluorescence confocal microscope. Calcofluor stains the live cells very rapidly and it does not appear to affect the cell morphology as the cells were intact and grew in coccal chains. The images were obtained with a high signal to noise ratio but without saturation effect. These images were then analyzed to understand whether calcofluor can be used to differentiate the HA producing cells from the non-producers.

### GJP2 cells exhibit higher HA expression

We first analyzed whether there is a difference between the expression levels of the GJP2 and the host (normal) phenotype. An example of the microscopy images in each case is shown in **Figure 5**, while the distribution of the expression values of individual cells is shown in **Figure 6**. We found that the expression of the GJP2 phenotype was consistently at a higher level. Specifically, we confirmed that the medians (0.0747 versus 0.387, fold-change of ~1.93) differ significantly between the GJP2 and Host phenotypes (rank-sum test, p-value < 1.95 × 10^−7^), in favor of higher median in the GJP2 phenotype, and that the differences between the GJP2 and Host expression are significantly larger than the differences between the images of each phenotype (F-test, p-value < 10^−8^).

**Figure 5:**
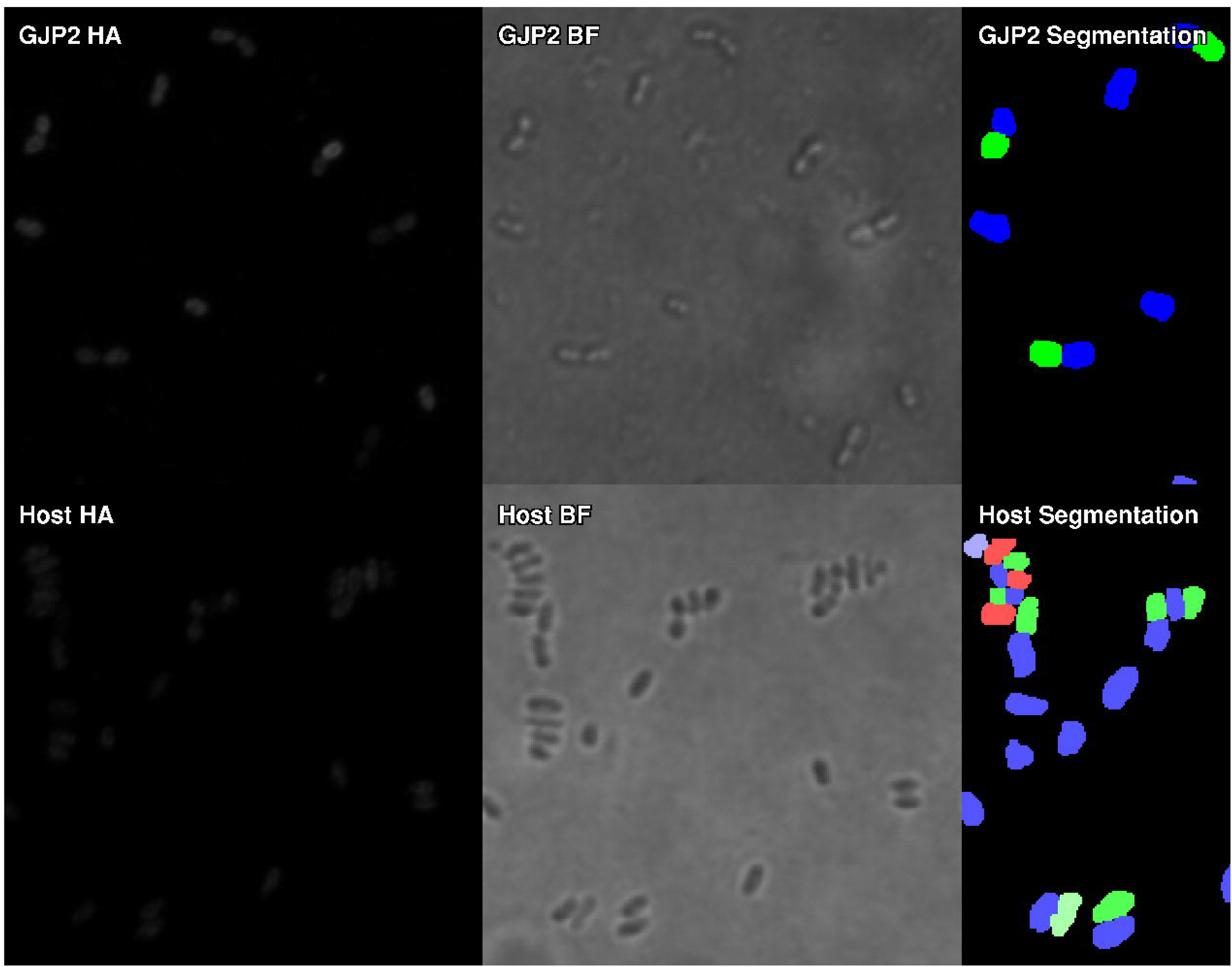
Example of microscopy images. The top rows show a region for one of the GJP2 cell images, while the bottom row shows that of the Host (normal) phenotype. The leftmost column shows the fluorescent HA signal, the middle column the bright field images used for cell segmentation and the rightmost column the segmented cells (using primary red, green, and blue colors with the saturation indicating the cell classification to dark, low, and high expression).

**Figure 6:**
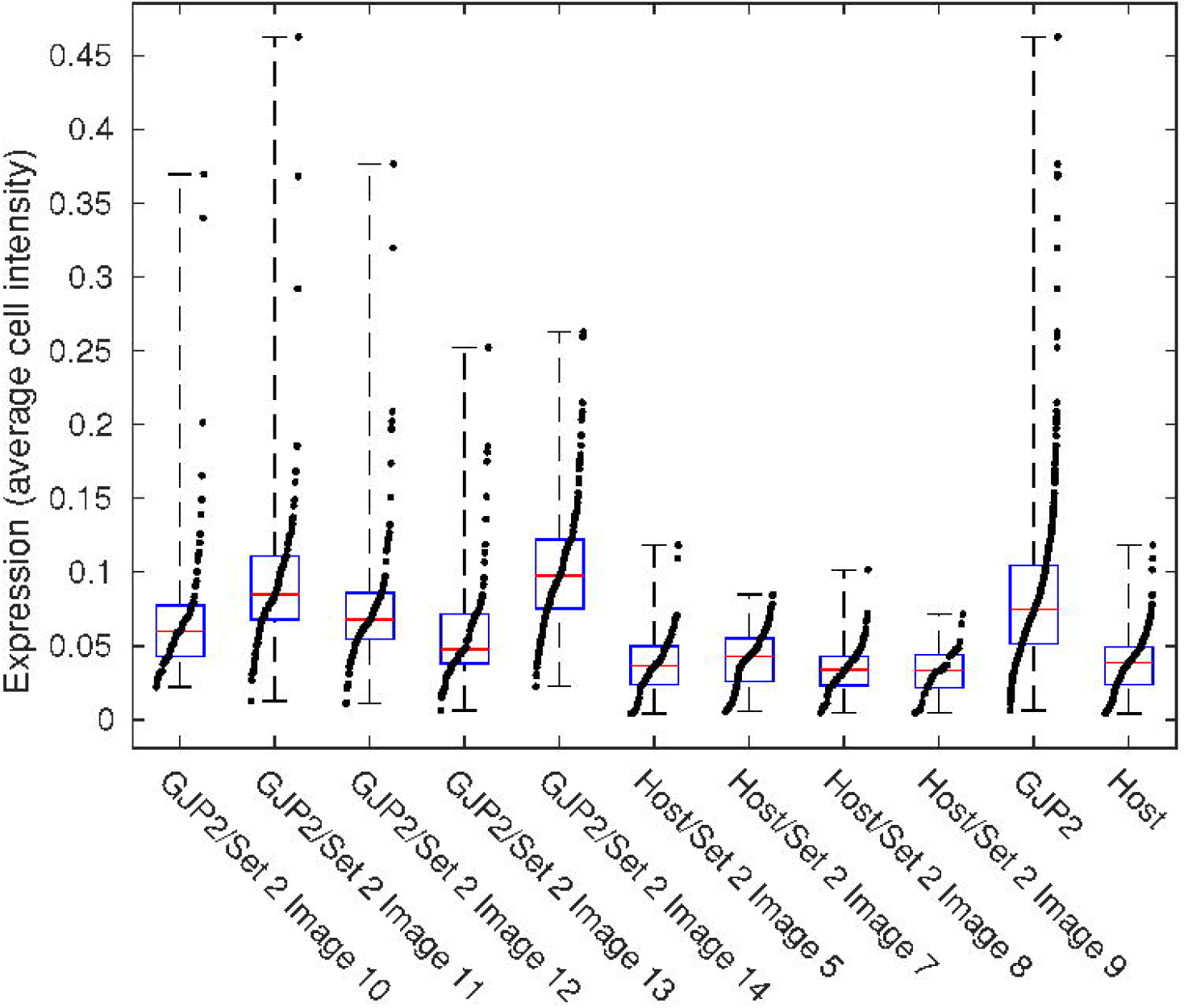
Expression of the HA marker in the GJP2 and host (normal) phenotypes from the fluorescence microscopy. Cellwise distribution of expression levels for each individual and image and groupwise aggregates. Red line denotes median, the box 25% to 75%, whiskers all data, and the dots individual cells jittered by their rank.

### GJP2 facilitates a high expression phenotype

We further investigated whether the higher expression of the GJP2 cells were due to higher expression level in general, or due to increased cellular heterogeneity. For this, we classified each cell as dark, low, or high expression by clustering the expression data into three ordered categories (see Methods). We found that the data are best explained by classifiying the GJP2 cells into high and low expression phenotypes while classifying the Host cells into low and dark classes (F-test against independent clusterings, p-value > 0.80). A scatterplot of the expression as function of the cell area and the classwise distributions of the expression values is shown in **Figure 7**. We also found the expression to be mostly independent of the cell size (area).

**Figure 7:**
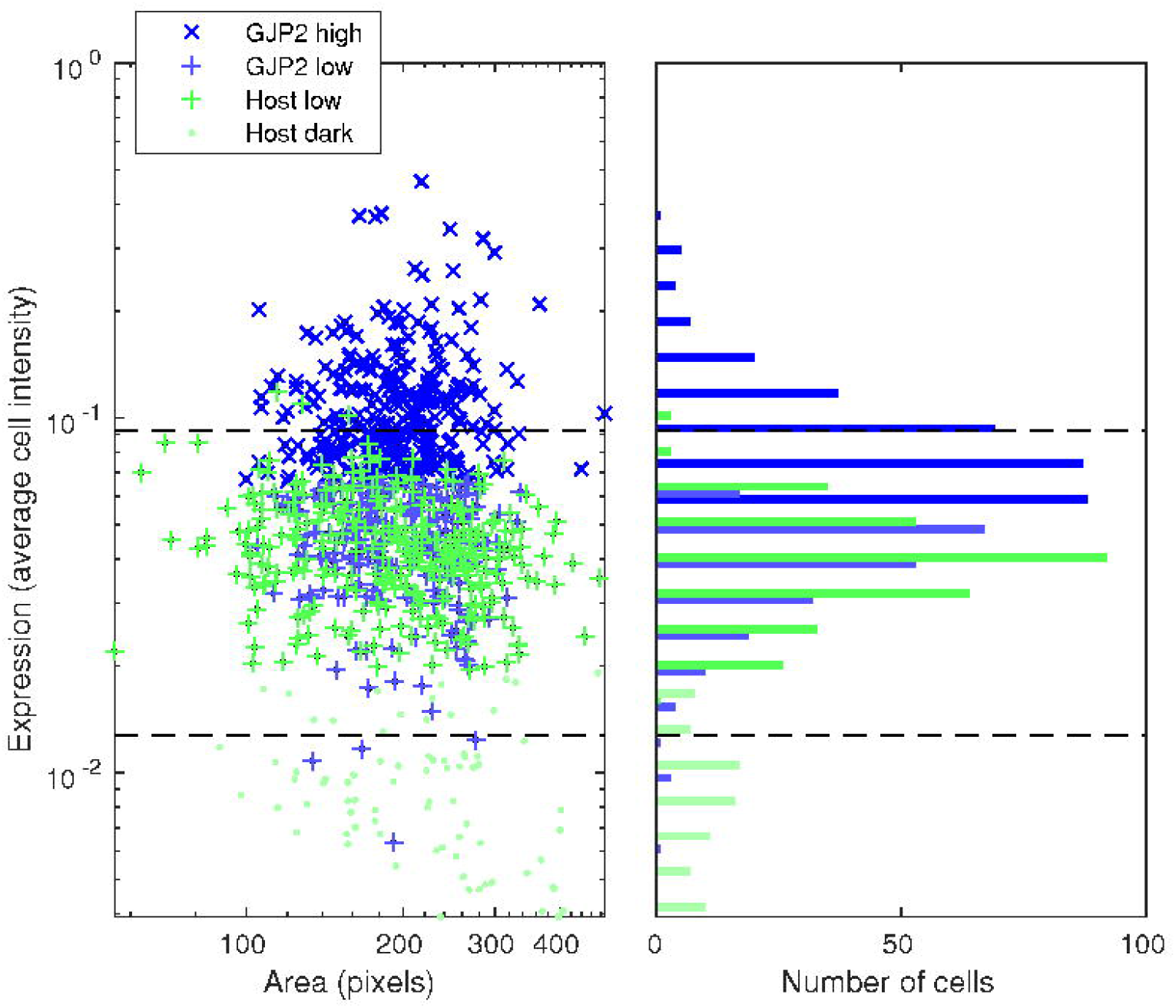
Clustered expression of the HA marker from the fluorescence microscopy. Left panel: expression versus cell area for cells clustered into dark, low, and high expression phenotypes. Right panel: distribution of expression levels for each cluster. The dashed lines indicate the cluster thresholds.

We found that each GJP2 phenotype image consistently features ~40% high expression cells and 60% low expression cells, while each Host image contains ~80% low expression cells and 20% dark cells. The dark cells feature expression level lower than ~0.0127 (median of ~0.0090) and the low expression cells more than that but lower than ~0.0921 (median of ~0.0438; see **Figure 7** and compare with **Figure 5**), while the high expression cells feature a median of ~0.0951 and fold-change of ~2.17 compared with the low expression phenotype. The exact numbers are tabulated in **Table 4**. This suggests that the exerted phenotypic changes are more drastic than those suggested by the population averages but not all the cells adopt this high expression phenotype.

Based on these results we concluded that the higher average expression harbored by the GJP2 phenotype is due to the emergence of a high HA expression phenotype, parallel to the normal phenotype, which is not present in the normal population. A source of suboptimality remains in the GJP2 population in terms of HA production, due to not all cells adopting the high expression phenotype.

## DISCUSSION

We developed an *in vivo* monitoring method based on calcofluor staining to differentiate HA producing *L. lactis* cells from non-producers. From the image analysis results, it is possible to conclude that our methodology can differentiate the phenotypes of *L. lactis* population with respect to presence of HA capsule. The host cells are either dark or having very low intensity, whereas the GJP2 population, which are harboring the genes for HA production and induced for the same, lacks dark cells found in the host phenotype. On the contrary, they feature high and low expression phenotypes, suggesting that this method is might be capable of differentiating the cells also based on the amount of HA present.

The distribution of phenotypes in the GJP2 cells implies that a source of suboptimality in HA production is a consequence of a notable fraction of low expression phenotypes in the population. Our results suggest that if this suboptimality could be removed, i.e. the population were entirely of the high expression phenotype, the HA production of our *L. lactis* strain could be improved to ~1.2 g/L. Also, the existence of heterogenous expression levels in the population might be a potential factor for high polydispersity (1.5) of HA.

Overall, this study contributes in optimizing genetic factors and process parameters more efficiently as it facilitates rapid *in vivo* screening of the population from the bioreactor samples. In the future, we believe that this method might help researchers to assess the impact of different intrinsic and extrinsic factors of gene expression on improving the high expression phenotypes in the population. Such studies might provide insights on improving the yield of HA in engineered cells and also reducing polydispersity by means of achieving a more homogenous population.

## Supporting information

Image analysis data

## Authors’ contributions

A-BM conceived the study. A-BM and AK jointly designed the study. A-BM performed Microscopy and AH performed image analysis. A-BM and VD jointly performed the experimental research. A-BM and AH analyzed data. A-BM, AH and GJ wrote the manuscript. All authors read and approved the manuscript.

## Acknowledgements

We thank Indian Institute of Technology Madras for providing infrastructure to carry out this work and Department of Biotechnology, Government of India for providing Fellowships to A-BM, VD and MHRD for AK. Further, we thank TRPVB, Madhvaram for the fluorescence confocal microscopy facility.

## Funding information

This work was supported by the Department of Biotechnology (Ministry of Science and Technology, Govt. of India) Project No.BT/PR13815/BBE/117/61/2015. AH is funded by Academy of Finland, grant No. 322927.

## Compliance with ethical standards

### Conflict of interest

The authors declare that they have no conflict of interest.

### Ethical approval

This article does not contain any studies with human participants or animals performed by any of the author.

### Consent to participate

Consent obtained from individual authors

### Consent for publication

Not applicable

### Availability of data and material (data transparency)

The measurement data are provided in Supplementary File 1.

### Code availability

Available upon request.

## Tables

**Table 1:** mCherry reporter production as quantified by the fluorometer as a function of the inducer (nisin) concentration. The data contains measurements from three replicates, their mean and standard error of the mean.

**Table 2:** HA yield in bioreactor studies using the CTAB analysis. The data contains measurements from two replicates, their mean and standard error of the mean.

**Table 3:** Molecular weight of the HA from size exclusion chomatography measurements.

**Table 4:** Groupwise expression statistics for each image. The table shows the number of dark, low, and high expression cells and their median (Med) and mean absolute deviation (Mad).

## Supplementary files

**Supplementary file 1**: Raw data at single-cell level after image analysis. The table contains the image, arbitrary cell (segment) IDs, summed intensity within the cell, the cell area (pixels), and the assigned expression class of the cell based on its average intensity.

---

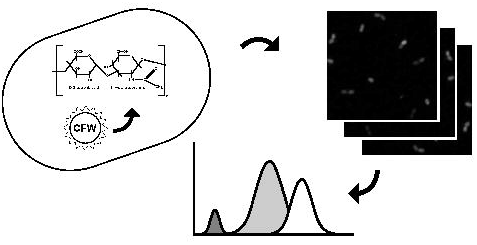

## Notes

### Competing Interest Statement

The authors have declared no competing interest.

